# Antiviral activity of the non-steroidal anti-inflammatory drug indomethacin against Respiratory Syncytial Virus

**DOI:** 10.1101/2025.04.28.650844

**Authors:** Caterina Tramontozzi, Anna Riccio, Simone La Frazia, Valentina Svicher, M. Gabriella Santoro

## Abstract

Human respiratory syncytial virus (RSV) is a ubiquitous pathogen belonging to the *Pneumoviridae* family. The enveloped virion contains a single-stranded, negative-sense RNA genome comprising 10 genes encoding 11 proteins. Infection is critically dependent on the RSV fusion (F)-protein, which mediates fusion between the viral envelope and host cells, as well as fusion of infected cells with adjacent cells, resulting in syncytium formation. RSV is the leading cause of hospitalization in children younger than 5-years of age with acute respiratory infections, posing a high risk of bronchiolitis and severe pneumonia particularly in infants under 6-months of age. RSV is also recognized to cause a substantial health burden in older adults. Virus-triggered exacerbated inflammation is associated with severe RSV disease outcomes. Although several F-targeting vaccines and monoclonal antibodies are now available for prevention of RSV disease in infants and older adults in several countries, there is no effective specific drug treatment for RSV. Herein we show that the non-steroidal anti-inflammatory drug indomethacin has a remarkable antiviral activity against RSV, causing a decrease in infectious virus production in different types of human respiratory cells at low micromolar concentrations. Whereas the mechanisms underlying the antiviral activity against RSV remain to be elucidated, we found that indomethacin does not affect RSV adsorption or entry into host cells, but acts at postentry level interfering with F-protein expression. The results suggest that indomethacin, possessing both anti-inflammatory properties and a direct antiviral activity against RSV, may represent an useful tool for the treatment of respiratory syncytial virus infections.

## Introduction

Respiratory syncytial virus (RSV) is a filamentous enveloped RNA virus belonging to the *Pneumoviridae* family. The virion contains a single-stranded, unsegmented, negative-sense 15.2 kb RNA genome comprising 10 genes encoding 11 proteins, surrounded by a lipid bilayer displaying the fusion (F), attachment (G) and, to a lower extent, small hydrophobic (SH) proteins (Langedijk and Bont, 2023). Infection is critically dependent on the RSV F-protein, which mediates fusion between the viral envelope and airway epithelial cells, as well as fusion of infected cells with adjacent cells, resulting in multinucleate syncytium formation (Battles and McLellan, 2019).

RSV is a ubiquitous pathogen generally transmitted through close contact and aerosolized droplets, and is classified into two major subtypes, A and B, both of which are known to circulate at varying levels depending on the RSV season (Nuttens et al., 2024). Whereas in most people it causes mild respiratory illness, RSV is the leading cause of hospitalization in children younger than 5 years of age with acute respiratory tract infections (ARI), posing a high risk of bronchiolitis and severe pneumonia particularly in infants under 6 months of age (Shi et al., 2017); in 2019 alone, RSV was estimated to be responsible for over 100,000 deaths globally among children under five years of age (Li et al., 2022). In addition to young children, RSV is recognized to cause a substantial health burden in older adults, especially among those with comorbidities and risk factors that often lead to hospitalization and severe outcomes (Falsey and Walsh, 2000), provoking global interest in the development of effective anti-RSV interventions.

Despite continuous attempts to develop vaccines since RSV discovery, until recently no effective vaccine was available, and RSV prevention among infants has relied on palivizumab, a short-term monoclonal antibody administered in five monthly doses to high-risk infants (Gutfraind et al., 2015). Two recombinant prefusion F protein-based (RSVpreF) vaccines (Abrysvo and Arexvy) are now available for the prevention of RSV disease in individuals 60 years of age and older. Abrysvo was recently approved by FDA and other regulatory agencies for active immunization of pregnant women for the prevention of RSV-caused ARI in infants from birth through 6 months of age (Du et al., 2025; Hong-Nguyen et al., 2025). In addition, an mRNA-based RSVPreF vaccine also received FDA-approval (Mullard, 2024). Alongside these vaccines, a long-acting monoclonal antibody, nirsevimab, has been recently approved for immunization of both preterm and term infants against RSV-related ARI in several countries (Drysdale et al., 2023). Monoclonal antibodies prophylaxis, however, is costly and cannot be broadly applied.

While some F-protein inhibitors are being investigated in randomized controlled trials [RV521 (sisunatovir) and AK0529 (ziresovir)], treatment options for RSV infection are limited to symptomatic therapy; the antiviral ribavirin, that shows *in-vitro* efficacy against RSV (Hruska et al., 1980), was found to have limited efficacy in patients and is no longer recommended (Sake et al., 2024). Therefore, the need for new therapeutic options continues to be relevant.

The non-steroidal anti-inflammatory drug indomethacin (INDO) is widely used in the clinic for its potent anti-inflammatory and analgesic properties (Vane and Botting, 1998; Rouzer and Marnett, 2009). INDO acts as a nonselective inhibitor of the enzymes essential for the synthesis of prostaglandins and thromboxane, cyclooxygenase-1 and -2 (COX-1 and COX-2), where COX-1 is the constitutively expressed isoform, and COX-2 is primarily an inducible enzyme, whose expression is activated in response to several stimuli, such as cytokines, endotoxin, hyperthermia and viral infection, including RSV infection (Rouzer and Marnett, 2009; Rossi et al., 2012; Richardson et al., 2005).

In addition to the anti-inflammatory action, INDO was shown to also possess antiviral properties. Since the initial discovery, *in-vitro* and *in-vivo* studies have confirmed the antiviral activity of indomethacin against several DNA and RNA viral pathogens, including herpesvirus (Zhu et al., 2002), hepatitis B (Bahrami et al., 2005), rotavirus (Rossen et al., 2004) and human immunodeficiency virus (Bourinbaiar et al., 1995). We have previously shown that INDO is also effective against both negative-sense RNA viruses (Amici et al., 2015) and positive-sense RNA viruses, such as canine and human coronaviruses, including SARS-CoV-1 (Amici et al., 2006; Tramontozzi et al., 2024). Antiviral effects of INDO have also been recently shown against SARS-CoV-2 in *in-vitro* models (Wang et al., 2023), as well as in clinical trials for the treatment of COVID-19 patients (Ravichandran et al., 2022).

We now show that INDO also effectively inhibits the replication of RSV in different types of human respiratory cells, acting at post-entry level.

## Results and Discussion

To investigate the effect of indomethacin (INDO, Fig. 1C) on RSV replication, HEp-2 cells, an immortalized human respiratory epithelial cell line used to study RSV for decades, were infected with human RSV strain A2 at a multiplicity of infection (MOI) of 0.01 TCID_50_/cell and, after the 2h adsorption period, were treated with different concentrations of INDO. Controls received equal amounts of vehicle. At 48h post infection (p.i.), viral titers were determined in the supernatant of infected cells by TCID_50_ infectivity assay (Fig. 1A, B) as described (Santoro et al., 1988). In parallel, the effect of INDO on the viability of mock-infected HEp-2 cells was determined by MTT assay (Piacentini et al., 2023) (Fig. 1A). INDO was found to have a remarkable antiviral activity, reducing virus yield dose-dependently with an IC_50_ of 4µM, LD_50_ above 400µM and selectivity index >100 (Fig. 1A-C); a greater than 98% decrease in virus yield was detected at non cytotoxic concentrations. Next, HEp-2 cells were infected with RSV at different MOI (0.01, 0.1 or 1 TCID_50_/cell), and treated with INDO (10-100µM) or vehicle after the adsorption period; virus yield was determined at 48h p.i. as described above. The results, shown in Fig. 1D, demonstrate that INDO was effective under all conditions used, indicating that INDO antiviral activity is not affected by the number of RSV infectious particles. Moreover, at the concentration of 100µM, INDO was found to partially protect HEp-2 cells from the cytopathic effect caused by RSV infection (1 TCID_50_/cell) up to 24h p.i. (Fig. 1E).

**Figure 1.**
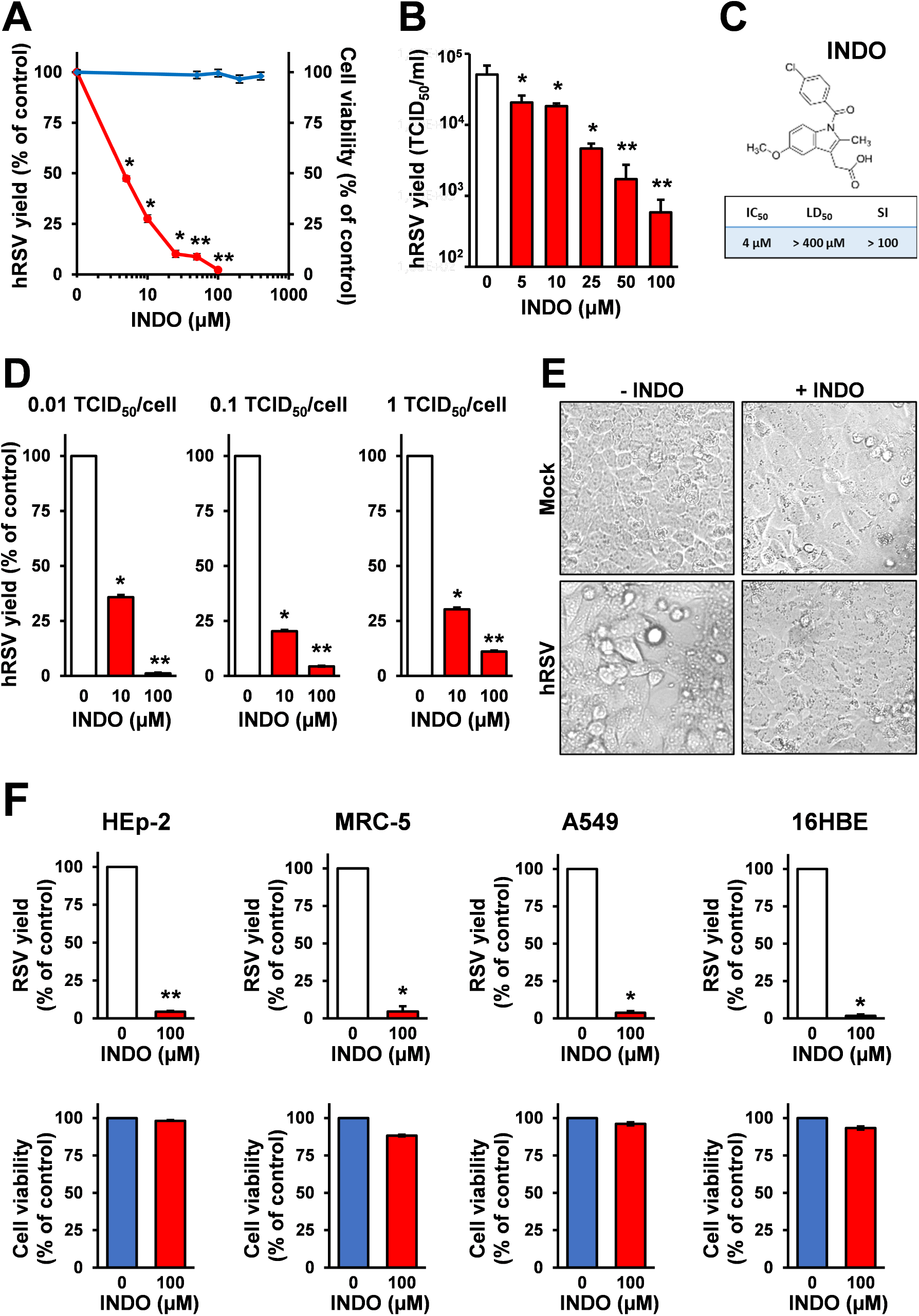
Indomethacin inhibits RSV replication in different types of human cells. (**A, B**) Human HEp-2 cell monolayers infected with RSV at an MOI of 0.01 TCID_50_/cell were treated with different concentrations of INDO or vehicle (0) after the adsorption period. Virus yield (red line) was determined at 48h p.i. by TCID_50_ infectivity assay. Data, expressed as percentage of control (A), represent the mean ± S.D. of triplicate samples. * = p < 0.05; ** = p < 0.01; ANOVA test. Data, expressed as TCID_50_/ml (B), represent the mean ± S.D. of duplicate samples from a representative experiment of three with similar results. * = p < 0.05; ** = p < 0.01; ANOVA test. In parallel, cell viability (A, blue line) was determined by MTT assay in mock-infected cells. Absorbance (O.D.) of converted dye was measured at λ = 570 nm. Data, expressed as percentage of control, represent the mean ± S.D. of triplicate samples. (**C**) Structure of indomethacin (INDO, upper panel). Indomethacin IC_50_ and LD_50_ in µM were calculated using Prism 8.0 software; selectivity index (SI) is indicated (C, lower panel). (**D**) HEp-2 cell monolayers infected with RSV at an MOI of 0.01, 0.1 or 1 TCID_50_/cell were treated with INDO (10 or 100 µM) or vehicle (0) after the adsorption period. Virus yield was determined at 48h p.i. by TCID_50_ infectivity assay. Data, expressed as percentage of control, represent the mean ± S.D. of triplicate samples from a representative experiment of two with similar results. * = p < 0.05; ** = p < 0.01; ANOVA test. (**E**) Cytoprotective effect of INDO (100µM) in HEp-2 cells infected with RSV (1 TCID_50_/cell) at 24h p.i. (magnification: 100×). **(F)** HEp-2, MRC-5, A549 and 16HBE cells were mock-infected or infected with RSV (0.1 TCID_50_/cell) and treated with INDO (100µM) or vehicle (0) after the adsorption period. Virus yield was determined at 48h p.i. by TCID_50_ infectivity assay (top panels). Data, expressed as percentage of control, represent the mean ± S.D. of triplicate samples. * = p < 0.05; ** = p < 0.01; Student’s *t*-test. In parallel, cell viability of mock-infected HEp-2, MRC-5, A549 and 16HBE cells treated with 100µM INDO or vehicle (0) was determined by MTT assay at 48h after treatment (bottom panels). Data, expressed as percentage of control, represent the mean ± S.D. of triplicate samples.

To determine whether INDO antiviral activity is dependent on the cell type, the effect of INDO on HEp-2 cells was compared to the drug activity in different types of human respiratory cells including normal lung MRC-5 fibroblasts, alveolar type II-like epithelial A549 cells, and human bronchial epithelial 16HBE cells. HEp-2, MRC-5, A549 and 16HBE cells were infected with RSV (0.1 TCID_50_/cell) and treated with 100µM INDO after the adsorption period. At 48h p.i. virus yield was determined in the supernatant of infected cells by TCID_50_ infectivity assay; in parallel, the effect of INDO on the viability of mock-infected cells was determined by MTT assay. As shown in Fig. 1F, INDO treatment was effective in inhibiting RSV replication in all types of human respiratory cells tested at non cytotoxic concentrations.

Next, to investigate whether INDO affects RSV receptor-binding or entry into host cells, HEp-2 cells were infected with RSV (0.01 TCID_50_/cell) and treated with 100µM INDO 6h before infection, or only during the 2h adsorption period, after which time the drug was removed; alternatively, RSV-infected cells were treated immediately after the adsorption period in the absence or the presence of 6h pretreatment. At 48h p.i. virus yield was determined as described above. As shown in Fig. 2A, INDO pretreatment did not affect RSV replication when the drug was removed before infection; treatment with the drug only during the virus adsorption period was also ineffective. On the other hand, treatment started after infection remarkably inhibited RSV infectious particles production, even in the absence of drug pretreatment, indicating that INDO acts at post-entry level.

**Figure 2.**
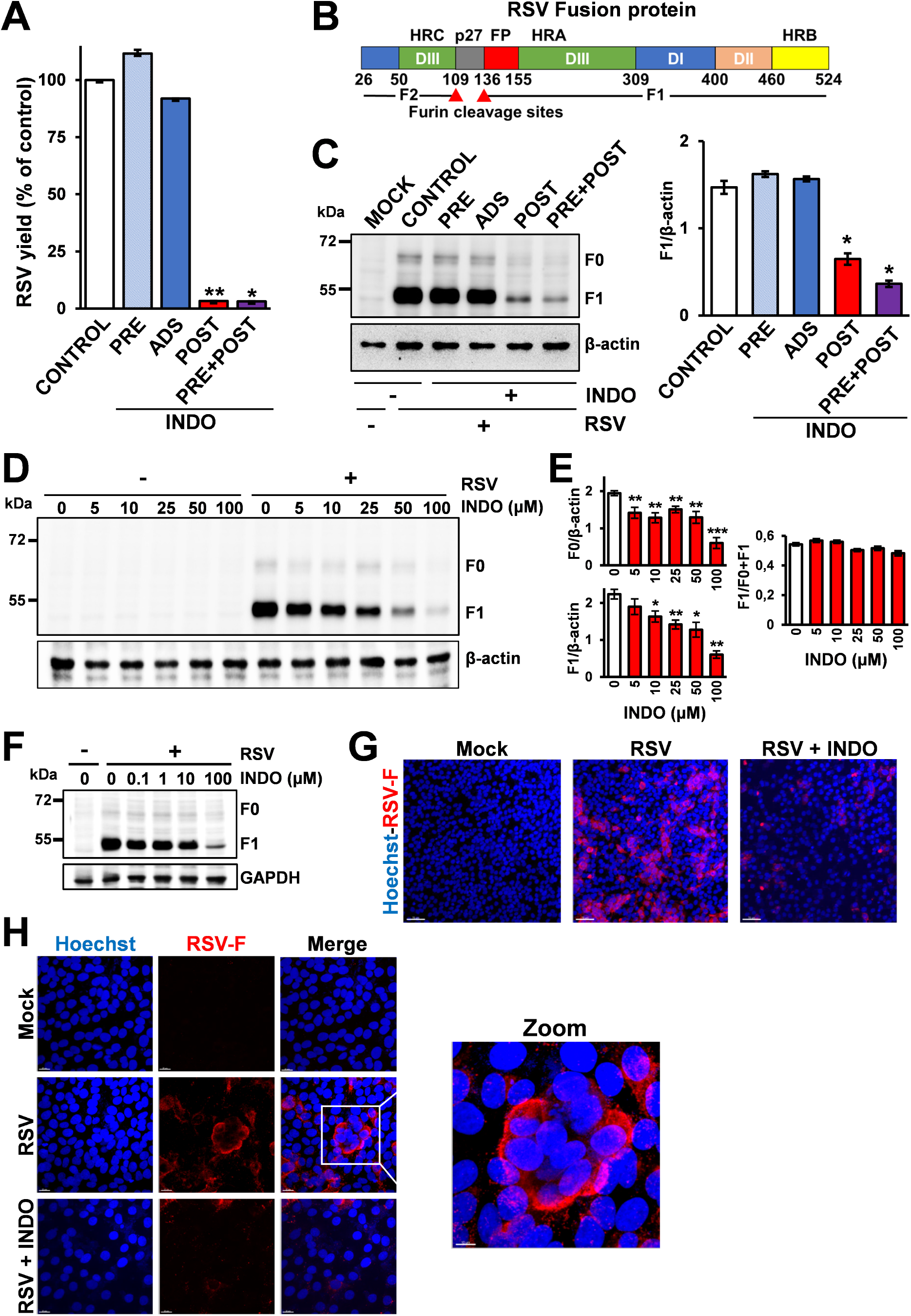
Indomethacin treatment decreases RSV F-protein levels, acting at post-entry level. **(A)** HEp-2 cells infected with RSV (0.01 TCID_50_/cell) were treated with 100µM INDO (+) or vehicle (-) at 6h before infection (PRE), only during the 2h adsorption period (ADS) or immediately after the adsorption period in the absence (POST) or the presence of 6h pretreatment (PRE+POST). Virus yield was determined at 48h p.i. by TCID_50_ infectivity assay. Data, expressed as percentage of control, represent the mean ± S.D. of duplicate samples from a representative experiment of two with similar results. * = p < 0.05; ** = p < 0.01; ANOVA test. (**B**) Schematic representation of RSV Fusion (F) protein structure. Residue numbers 26-524 are indicated. Protein domains are illustrated: DI-III, domains I-III; HRA, -B, and -C, heptad repeats A, B, and C; p27, excised peptide; FP, fusion peptide. The furin cleavage sites (red arrowheads) are shown. **(C)** Immunoblot analysis of RSV F-protein in HEp-2 cells treated as in (A) at 48h p.i.; the uncleaved F precursor (F0) and the F1 subunit are indicated (left panels). F0 and F1 levels were determined by densitometric analysis using ImageJ software, normalized to the loading control in the same sample, and expressed as arbitrary units (right panels). Error bars indicate the mean ± S.D. of duplicate samples from a representative experiment of two with similar results. * = p < 0.05; ANOVA test. (**D-F**) HEp-2 cell monolayers mock-infected (-) or infected (+) with RSV at an MOI of 0.01 TCID_50_/cell (D,E) or 0.1 TCID_50_/cell (F) were treated with different concentrations of INDO or vehicle (0) after the adsorption period. At 48h p.i., levels of the viral fusion protein were determined by immunoblot analysis as in (C). Relative amounts of the F0 precursor and the F1 subunit were determined by densitometric analysis, normalized to β-actin in the same sample, and expressed as arbitrary units (E, left panels). F1/F0+F1 ratio for each sample is shown (E, right panel). Error bars indicate the mean ± S.D. of duplicate samples from a representative experiment of two with similar results. * = p < 0.05; ** = p < 0.01; *** = p < 0.001; Student’s *t*-test. (**G**) Immunofluorescence analysis of RSV-F protein (red) in mock-infected and RSV-infected (1 TCID_50_/cell) HEp-2 cells treated with INDO (100µM) or vehicle for 24h. Nuclei are stained with Hoechst (blue). Merge images are shown. Scale bar, 70 µm. (**H**) Confocal images of RSV-F protein (red) in HEp-2 cells treated as in (G). Nuclei are stained with Hoechst (blue). Merge images are shown. Scale bar, 20 µm (zoom: 10 µm).

We have previously shown that INDO causes a dysregulation of viral structural protein expression in a different model of negative-strand RNA virus infection, Vesicular Stomatitis virus (VSV), acting at the translational level, by rapidly inducing the phosphorylation of the alpha-subunit of eukaryotic initiation factor-2 (eIF2α) via activation of the double-stranded RNA-activated protein kinase (PKR) (Amici et al., 2015; Brunelli et al., 2012). Among the different RSV structural proteins, the Fusion-protein is one of the most studied due to its critical role in mediating fusion and entry into host cells, as well as being responsible, together with the attachment glycoprotein G, for inducing neutralizing antibodies *in-vivo* (Swanson et al., 2011).

The RSV F-protein is a class I viral fusion glycoprotein that is synthesized as a 574-amino-acid inactive precursor called F0 that shares structural similarities with the fusion glycoproteins of other members of the *Pneumoviridae* and *Paramyxoviridae* (Battles and McLellan, 2019). In order to become fusion competent, F-protein must be cleaved by furin-like host proteases at two polybasic sites that are separated by 27 amino acids (p27) (Fig. 2B). Cleavage, which occurs in the *trans-*Golgi network during transport to the plasma membrane, generates two subunits, the amino-terminal F2-subunit and carboxy-terminal F1-subunit. F1 is anchored in the viral membrane via a transmembrane region and harbors a N-terminal hydrophobic fusion peptide (FP, Fig. 2B) responsible for cellular membrane insertion; residues in the F2-subunit contribute to F-protein fusogenicity and possibly to RSV species-specificity (Gilman et al., 2019). The two subunits are covalently linked via two disulfide bonds and associate to form the mature, fusion-competent trimeric F-protein form. As indicated above, when expressed on the plasma membrane of infected cells RSV F-protein also mediates cell-to-cell fusion, resulting in multinucleate syncytium formation.

The effect of INDO on the expression of the RSV F-glycoprotein was therefore investigated. First, HEp-2 cells infected with RSV (0.01 TCID_50_/cell) were treated with 100µM INDO 6h before infection, only during the 2h adsorption period, or after the adsorption period, as described above, and at 48h p.i. F-protein levels in infected cells were determined in whole-cell extracts (WCE) by immunoblot analysis (Coccia et al., 2017) using specific antibodies recognizing the F0-precursor and the F1-subunit, followed by scanning densitometry, as previously described (Riccio et al., 2022). As shown in Fig. 2C, INDO pretreatment or treatment only during the virus adsorption period did not affect F-protein levels when the drug was removed before infection, whereas treatment started after infection caused a remarkable decrease in F-protein expression even in the absence of drug pretreatment, confirming that INDO acts at post-entry level.

Next, HEp-2 cells were mock-infected or infected with RSV at an MOI of 0.01 (Fig. 2D) or 0.1 (Fig. 2F) TCID_50_/cell and treated with different concentrations of INDO after the adsorption period. At 48h p.i., F-protein levels were analyzed in WCE by immunoblot analysis. INDO treatment caused a dose-dependent decrease in F-protein levels; the F0-precursor and the F1-subunit were found to be similarly affected, indicating that INDO did not significantly alter F0 cleavage (Fig. 2E). Decreased levels of F-protein were also detected by immunofluorescence in RSV-infected (1 TCID_50_/cell) HEp-2 cells treated with 100µM INDO after the adsorption period for 24h (Fig. 2G). A decrease in RSV F-mediated syncytia formation was also observed in INDO-treated cells, as a consequence of lower F-protein levels (Fig. 2H).

Taken together, the results show that indomethacin at low micromolar concentrations inhibits RSV replication in *in-vitro* models, acting at post-entry level and causing a decrease in F-protein expression. The mechanisms underlying the antiviral activity of INDO against RSV remain to be elucidated. The ability of INDO to inhibit the replication of several unrelated RNA and DNA viruses points out to a host-mediated antiviral mechanism, rather than an effect on virus-specific enzymes. Whereas INDO anti-inflammatory action is mediated by blocking COX-1 and COX-2 enzymatic activity (Rouzer and Marnett, 2009), during VSV and coronavirus infection the antiviral activity was found to be cyclooxygenase-independent and it could not be mimicked by other COX-1/2 inhibitors (Amici et al., 2015; Tramontozzi et al., 2024); moreover, it should be pointed out that INDO antiviral effects occur at concentrations higher than those needed for COX inhibition (10^−8^-10^−9^M). In the case of RSV, it is important to note that RSV infection induces COX-2 expression in human lung cells *in-vitro*, as well as *in-vivo* in the lungs of cotton rats, leading to increased prostaglandins production, an effect associated with RSV inflammatory pathogenesis (Richardson et al., 2005). Interestingly, administration of indomethacin significantly decreased lung pathology, including bronchiolitis, alveolitis, and interstitial pneumonitis, associated with RSV infection in the cotton rat; this effect was ascribed to the drug anti-inflammatory rather than antiviral action (Richardson et al., 2005). These observations, together with the anti-RSV activity of INDO in human respiratory cells described in the present report, suggest that indomethacin, possessing both anti-inflammatory properties and a direct antiviral activity against RSV, could be an useful tool for the treatment of respiratory syncytial virus infections.

## Materials and Methods

### Cell culture and treatments

Epithelial HEp-2 cells derived from human larynx carcinoma, human alveolar type II-like epithelial A549 cells, human normal lung MRC-5 fibroblasts and human bronchial epithelial 16HBE cells were obtained from American Type Culture Collection (ATCC; Manassas, VA, USA). Cells were grown at 37°C in a 5% CO_2_ atmosphere in RPMI-1640 (Euroclone; A549 and HEp-2), DMEM (Euroclone; 16HBE) and EMEM (ATCC; MRC-5) medium, supplemented with 10% fetal calf serum (FCS), 2 mM glutamine and antibiotics. Indomethacin (Sigma) was dissolved in ethanol stock solution (40 mM), and diluted in culture medium. The compound was added to infected cells after the virus adsorption period and maintained in the medium for the duration of the experiment, unless differently specified. Controls received equal amounts of vehicle, which did not affect cell viability or virus replication. Cell viability was determined by the 3-(4,5-dimethylthiazol-2-yl)-2,5-diphenyltetrazolium bromide (MTT) to MTT formazan conversion assay (Sigma-Aldrich), as described (Piacentini et al., 2023). The 50% lethal dose (LD_50_) was calculated using Prism 8.0 software (Graph-Pad Software Inc.). Microscopical examination of mock-infected or virus-infected cells was performed using a ZEISS Axio Observer 7 inverted microscope with Apotome III and analyzed using ZEN 3.1 (blue edition) software.

### Virus production, infection and titration

Human respiratory syncytial virus A2 strain (RSV-A2), a kind gift from G. Toms, Newcastle University, was used for this study. For RSV-A2 production, confluent HEp-2 cell monolayers were washed twice with phosphate-buffered saline (PBS) and infected with RSV at a multiplicity of infection (MOI) of 0.1 TCID_50_/cell (50% tissue culture infectious dose)/cell. At 72h p.i. the virus was harvested by scraping the cells and collecting them together with the culture supernatants. Cells and supernatants were centrifuged (5 min, 4,000 × g, 4°C) and freeze-thawed 3 times; after vortexing, cell debris and supernatant were combined, aliquoted, and stored in liquid nitrogen as described (Sun and Lòpez, 2016). For RSV infection, confluent HEp-2, MRC-5, A549 and 16HBE cell monolayers were infected with RSV for 2 hours at 37°C at different MOI (0.01, 0.1 or 1 TCID_50_/cell). After the adsorption period, the viral inoculum was removed, and cell monolayers were washed three times with PBS and incubated at 37°C in medium containing 2% FCS. The virus was harvested as described above. Virus yield was determined at different times after infection by TCID_50_ infectivity assay, as described previously (Santoro et al., 1988). The 50% inhibitory concentration (IC_50_) of the compound tested was calculated using Prism 8.0 software.

### Protein Analysis and Western blot

For analysis of proteins, whole-cell extracts (WCE) were prepared after lysis in High Salt Buffer (HSB) as described (Santoro et al., 1982). Briefly, cells were washed twice with ice-cold PBS and then lysed in HSB (40 µl). For Western blot analysis, cell extracts (15 µg/sample) were separated by SDS-PAGE under reducing conditions and blotted to a nitrocellulose membrane. After blocking with 5% skim milk solution, membranes were incubated with the selected antibodies followed by decoration with peroxidase-labeled anti-rabbit or anti-mouse IgG. Primary and secondary antibodies used are listed in Table 1. Quantitative evaluation of proteins was determined as described (Santoro et al., 1989). All results shown are representative of at least three independent experiments.

**Table 1.** Antibodies used. (**M**): Monoclonal; (**P**): Polyclonal.

| Antibody | Source | Catalogue number |
| --- | --- | --- |
| β-Actin (P) | Sigma-Aldrich | A2066 |
| GAPDH (P) | Cusabio | CSB-PA00025A0Rb |
| RSV Fusion Protein (2F7) (M) | Novus Biologicals | NB110-37246 |
| Alexa Fluor<sup>TM</sup> 555 goat anti-mouse | Invitrogen | A21422 |
| Goat Anti-Mouse IgG (H+L), HRP | Jackson ImmunoResearch | 115-035-003 |
| Goat Anti-Rabbit IgG (H+L), HRP | Jackson ImmunoResearch | 111-035-003 |

### Immunofluorescence microscopy

HEp-2 cells grown in 8-well chamber slides (LabTek II, Nunch) were infected with RSV and, after the adsorption period, were treated with INDO or vehicle for 24h. Cells were fixed with 4% paraformaldehyde, permeabilized with 0.5% TritonX100-PBS (Riccio et al., 2022), and processed for immunofluorescence using selected antibodies followed by decoration with Alexa Fluor 555-conjugated antibodies (Molecular Probes, Invitrogen). Primary and secondary antibodies used are listed in Table 1. Nuclei were stained with Hoechst 33342 (Molecular Probes, Invitrogen). Images were captured using a ZEISS Axio Observer 7 inverted microscope with Apotome III and analyzed using ZEN 3.1 (blue edition) software. For confocal microscopy, images were acquired on Olympus FluoView FV-1000 confocal laser scanning system (Olympus America Inc., Center Valley, PA) and analyzed using Imaris (v6.2) software (Bitplane, Zurich, Switzerland). Images shown in all figures are representative of at least three random fields (scale-bars are indicated).

### Statistical analysis

Statistical analyses were performed using Prism 8.0 software (GraphPad Software). Comparisons between two groups were made using Student’s *t*-test; comparisons among groups were performed by one-way ANOVA with Bonferroni adjustments. *p* values ≤0.05 were considered significant. Data are expressed as the means ± standard deviations (S.D.) of results from duplicate or triplicate samples. Each experiment (in duplicate) was repeated at least twice.

## Author Contributions

CT, AR and SLF performed the study on the antiviral activity; CT and AR performed protein synthesis analysis. MGS designed the study. MGS, CT, AR and SLF wrote the manuscript. MGS, and VS revised the manuscript. All authors contributed to the interpretation of the data and approve the content of the manuscript.

## Acknowledgments

The authors thank Prof. G. Toms, University of Newcastle, Newcastle upon Tyne, UK, for kindly providing the human RSV-A2 virus. We also thank Dr. Elena Romano (University of Rome Tor Vergata) for assistance with confocal microscopy. Caterina Tramontozzi was enrolled in the PhD Program in Cellular and Molecular Biology, Department of Biology, University of Rome Tor Vergata, Rome, Italy.

## Funding

This research was supported by grants from the Italian Ministry of University and Scientific Research (PRIN project 2010PHT9NF-006) and from the “Fondo Intesa Sanpaolo” (project B/2023/018).

## Conflict of Interest statement

The authors declare no conflict of interest.

## Data availability statement

Data will be made available on reasonable request

## Notes

### Competing Interest Statement

The authors have declared no competing interest.

